# Negative frequency-dependent selection maintains partner quality variation in a keystone nutritional mutualism

**DOI:** 10.64898/2026.02.17.706392

**Authors:** Rebecca T. Doyle, Xingyuan Su, Christina Gallick, Meghan Blaszynski, Emily Perry, Karla Griesbaum, Omolabake O. Oyetayo, David Vereau Gorbitz, Jennifer A. Lau, Katy D. Heath

## Abstract

Mutualisms, interactions that benefit both partners, are critical for promoting biodiversity and ecosystem resiliency, yet are considered evolutionarily unstable and vulnerable to global change. Understanding how genetic variation in mutualisms is maintained is key to predicting their future persistence and explaining their long evolutionary history. While theory addresses factors maintaining variation in partner quality (the fitness benefits partners provide), few studies have experimentally tested these mechanisms. Here, we experimentally evolved multiple replicate populations of rhizobia, nitrogen-fixing mutualists of legumes, varying in partner quality under contrasting environments: nitrogen (N)-supplemented or N-free conditions, with or without host plants. After one year of rhizobial evolution, we quantified selection on partner quality across environments and evaluated resulting changes in mutualism traits and population-level genetic diversity. Strikingly, selection on partner quality was population-dependent: high-quality strains were favoured when initially rare but disfavoured when common, revealing negative-frequency dependent dynamics that can maintain variation. Although neither nitrogen-supplementation nor host presence directly imposed selection, both were critical for preserving genetic diversity in rhizobia populations – fuel for ongoing evolution. By demonstrating negative-frequency dependent selection in the legume-rhizobium mutualism, our study reveals a dynamic more akin to antagonistic interactions than traditionally assumed. This overlooked mechanism may be the key to explaining the ecological and evolutionary persistence of mutualisms under changing environments.

**Significance Statement:** Mutualisms are central to biodiversity, yet their stability is puzzling because the mechanisms thought to stabilize them tend to eliminate the genetic variation needed for continued adaptation, even though partners in nature show striking variation in quality. In an experimental evolution study of a keystone plant-microbe mutualism (legume–rhizobium symbiosis), we found that negative frequency-dependent selection allows high- and low-quality symbionts to coexist: high-quality strains are favoured only when rare. Environmental factors such as nitrogen addition or host presence did not alter which partners were selected, but they helped maintain genetic diversity needed for future adaptation. Our findings reveal an overlooked mechanism that stabilizes mutualisms and explain how these interactions can remain resilient under changing environmental conditions.

## Introduction

Mutualisms, or mutually beneficial interactions among species, are essential for promoting biodiversity and ecosystem resiliency (1). Despite that many well-known mutualisms are evolutionary ancient, including the arbuscule mycorrhizal fungus-plant mutualism estimated to have evolved more than 500 million years ago (2), these interactions are predicted to be evolutionarily unstable and susceptible to rapid global change (3–7). As a result, decades of theory have generated explanations for how mutualism can be maintained (6, 8–12). However, most, if not all, of these theoretical explanations for mutualism persistence result in the erosion of standing genetic variation in a key mutualism trait—partner quality, or the fitness benefit a partner confers to its interacting mutualist, often a host. Yet partner quality varies extensively in nature (13–16), calling into question what factors maintain such variation? Identifying the forces that maintain variation in mutualisms is critical for predicting their future persistence under global change and for resolving their paradoxically long evolutionary history (8).

For decades, mutualism theory has predicted that variation in partner quality should erode over time, either through the spread of low-quality cheaters that avoid the costs of cooperation or through strong host-mediated selection favouring high-quality partners, both of which should reduce diversity and potentially destabilize cooperation (8, 9, 17). Yet empirical and theoretical work increasingly challenges this view: high-quality partners often outcompete low-quality ones in experimental competitions (18–21), and partner fidelity feedbacks can tightly couple partner fitness to its cooperative value, limiting the success of cheaters (11, 12, 22, 23). At the same time, the forces that suppress cheating, including host choice and sanctions, are expected to impose directional or stabilizing selection that further depletes the genetic variation required for mutualism to evolve (8). These predictions contrast with the substantial partner-quality variation observed in nature, suggesting that additional mechanisms may sustain diversity. Possible processes include negative frequency-dependent selection, in which rare genotypes gain a fitness advantage (24); environmentally dependent selection, where abiotic or biotic context alters which partner phenotypes are favoured (25); and life-stage-specific trade-offs, such as low-quality partners performing better outside the host but poorly within it (26). Because empirical tests of these mechanisms remain limited, experiments that manipulate ecological conditions while tracking selection on mutualist traits are essential for understanding how variation in partner quality persists despite strong theoretical expectations of its erosion.

Given a rapidly changing environment, understanding the maintenance of genetic variation in partner quality is increasingly important, genetic variation being the prerequisite for mutualist responses to environmental change (27–29). For example, nitrogen (N)-based fertilizer supplementation in agriculture is a major anthropogenic contributor of nutrient loading, reliably boosting crop yields but at the cost of reducing soil microbial diversity, including the diversity of key mutualists like mycorrhizae and nitrogen-fixing bacteria (30, 31). The nutritional mutualism between legumes (e.g., beans, peas, alfalfa) and rhizobia is particularly vulnerable to nitrogen-supplementation because legumes rely less on symbiotic N fixation when external N is available (32). When soil N is limiting, legumes attract rhizobia via flavonoid signaling, triggering nodulation – the formation of root structures known as nodules that house rhizobia. However, sufficient N suppresses nodulation, reducing symbiotic investment and the fitness benefits of mutualism for the legume (33–36). Nitrogen also causes evolutionary declines in mutualism: multiple decades of nitrogen-supplementation in experimental field plots caused the evolution of lower quality rhizobia (37, 38). Because rhizobia alternate between a free-living soil stage and a symbiotic stage within host plants, they offer a unique opportunity for disentangling how selection in soil or host environments, independently or together, shapes population structure and the maintenance or loss of partner quality variation. Determining the type and direction of selection acting on rhizobium populations across distinct selective environments would thus be a critical step towards uncovering the factors important for driving partner quality variation and lead to more accurate predictions of whether this critical mutualism will persist under a rapidly changing environment.

Experimental evolution combined with resequencing (“evolve-and-resequence”) has served as a powerful tool for identifying the selective forces and genetic mechanisms driving microbial evolution (39). By tracking changes in strain frequencies and genotypes over time, this approach can reveal the roles of standing genetic variation, de novo mutation, and recombination in shaping evolutionary trajectories. While early applications focused on clonal, non-symbiotic microbes in static environments (e.g., Lenski’s *E. coli* experiment (40)), more recent studies have extended this framework to mutualisms, using metagenomics to track genotype frequencies in diverse symbiont populations exposed to distinct selective environments (20). Building upon these approaches by incorporating multiple genetically distinct rhizobial populations that vary in partner quality would allow for the simultaneous assessment of how environmental factors, including nitrogen availability and host presence, shape selection on partner quality, and whether selection is frequency dependent. Leveraging natural variation in mutualist traits, including partner quality, and exposing populations to ecologically relevant conditions provides a more nuanced understanding of the mechanisms that maintain, rather than erode, variation in mutualisms.

Here, we experimentally evolved multiple replicate populations of three rhizobium population types that differed in the initial frequency of high-quality strains: one where high-quality strains were rare (“Low-Quality” population), one where they were common (“High-Quality” population), and one with an intermediate mix (“Medium-Quality” population). We passaged replicate populations of each population type under two contrasting selective environments: nitrogen-supplemented (N+) versus nitrogen-limited (N-) conditions, as well as in the presence (P+) or absence (P-) of plant hosts across four plant growth cycles, each lasting ∼2 months, for a total of just under one year of rhizobial evolution (∼400 rhizobial generations). Each treatment combination was represented by 3-4 independently evolving replicate populations (see **Materials and Methods** for details), resulting in a total of 30 replicate populations across all selective environments. Our goals were to (1) quantify how selection on partner quality varied across population types and contrasting environments; (2) determine how populations evolving under these environments diverged in key mutualism traits, including population-level partner quality and host symbiotic investment; and (3) evaluate how selection influenced strain diversity, measured as the proportion of the original 28 strains recovered from each replicate population after experimental evolution.

Our experiment allowed us to test multiple, non-mutually exclusive predictions. If nitrogen supplementation drives directional selection favouring low-quality strains, as suggested by previous work (37, 38), then we would expect populations passaged under nitrogen-supplemented conditions to show negative directional selection on partner quality, a decline in average partner quality, and a reduction in genetic variation relative to those evolved under N-limiting conditions. Alternatively, if selection on partner quality is frequency-dependent, then the fitness advantage of high- versus low-quality strains should vary with their initial frequency in the population. Such frequency-dependent selection would result in divergent evolutionary outcomes across different starting populations, with greater variation in mean partner quality and final strain compositions. By evolving multiple genetically distinct rhizobial populations under contrasting selective environments, our approach allows us to simultaneously assess the roles of environmental context and initial population structure in shaping selection on partner quality. Our results not only reveal how selection operates in a keystone plant symbiont but also identify key ecological and evolutionary factors that help maintain variation in mutualist traits, offering new insight into how mutualisms evolve and persist over time.

## Results

### Negative frequency-dependent selection on rhizobium partner quality

To determine how selection acted on rhizobium populations across distinct selective environments, we used genome sequencing to track the frequency of each strain in each independently evolved replicate population (i.e., derived isolates; n = 362 total, 38 - 54 isolates per population type and selective environment), then used genotypic selection analysis (41, 42) to relate strain relative fitness to partner quality. In a global selection model (across all three population types), selection tended to favour strains of intermediate quality (i.e., significant non-linear quadratic (γ) term: -0.936 +/- 0.329 SE; F [1, 604] = 8.10, *P =* 0.005); however, selection operated differently across population types (i.e., significant partner quality x population type interaction term: F [2, 604] = 5.65, *P* = 0.004). These models were conducted at the strain level (n = 616 strain-level observations for analyses restricted to host-present treatments, and n = 392 observations for analyses restricted to the Medium-Quality population type; see **Materials and Methods**), and we found little evidence that selection depended on either nitrogen supplementation or host presence (i.e., non-significant partner quality x N-environment and partner quality x host presence interaction terms; **SI Appendix Table S1**).

Next, we conducted selection models within each population type (**SI Appendix Table S2**) using strain-level observations nested within replicate populations, and found that Low-Quality populations showed signatures of stabilizing selection on partner quality (quadratic selection term [γ]: -0.878 +/- 0.346 SE; F [1,221] = 6.44, *P* = 0.0119; top row, **Fig. 1A**). Significant positive directional selection differentials (*S*) were also detected (0.508 +/- 0.197; F [1,222] = 6.61, *P* = 0.011), and in response to this selection, the median partner quality of the evolved populations shifted towards significantly higher quality (t = 7.83, df = 7, *P* < 0.001; **Fig. 1B**). Similarly, Medium-Quality populations experienced stabilizing selection on partner quality (γ: -0.884 +/-0.181 SE; F [1,389] = 24, *P* < 0.001; middle row, **Fig. 1A**) with significant positive directional selection components (*S*: 0.266 +/- 0.103; F [1,390] = 6.7, *P* = 0.01), and the median partner quality of the evolved populations shifted slightly higher than the initial population (t = 5.77, df = 7, *P* < 0.001, **Fig. 1B**). In contrast, High-Quality populations experienced negative directional selection for partner quality (*S*: -0.675 +/- 0.155 SE; F [1,222] = 19, *P* < 0.001; bottom row, **Fig. 1A**), shifting evolved populations significantly towards lower partner quality (t = -5.18, df = 7, *P* = 0.001; coloured vertical lines in **Fig. 1B**). Consistent with these findings, permutation tests, which were conducted to account for replicate population-specific sampling effort (**SI Appendix Fig. S1**), showed that half (4/8) of the replicate populations in the Low-Quality treatment had higher-than-expected PQ (empirical *P* < 0.1) after evolution, none of the 14

**Figure 1.**
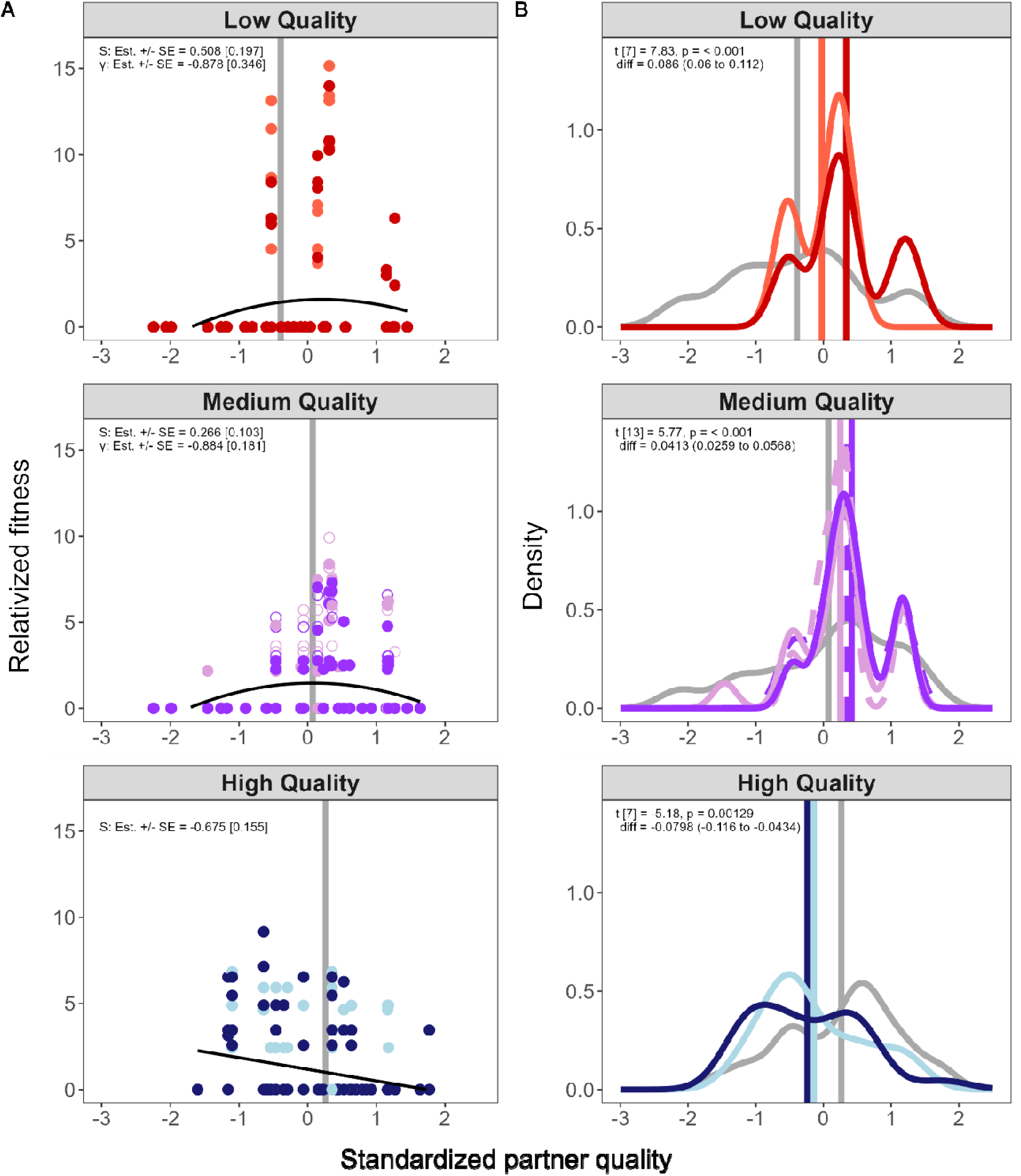
Selection on partner quality depends on the initial population type. **A** Selection analyses for strain means of standardized partner quality across all strains and relativized strain fitness within each selective environment. Vertical lines represent the median partner quality of each initial population. **B** Standardized partner quality distributions and medians (vertical lines) based on the strain composition of initial (gray) and experimentally evolved (colour) populations. The different shades within each panel show the two nitrogen environments (lighter = N-; darker = N+) while solid or dashed distributions show the two host plant environments (P+ or P-, respectively); selection did not differ between nitrogen or host plant treatments (see **SI Appendix Table S1**). Selection analyses were conducted at the strain level, with 28 strains quantified within each independently evolved replicate population (n = 30 replicate populations; 840 total strain observations).

Medium-Quality replicates deviated from their null distributions (*P* > 0.1), and half (4/8) of the replicate populations within the High Quality treatment had lower-than-expected PQ (empirical *P* < 0.1), indicating significant declines in partner quality in those populations.

Given that selection and evolutionary responses depended on the initial starting population conditions (Population Type) and the overall trend favouring strains of intermediate partner quality, we next tested for frequency-dependent selection by regressing the change in partner quality against each replicate population’s initial partner quality. This analysis, conducted at the population level (i.e., each replicate population was treated as an independent data point), revealed negative frequency-dependent selection: populations that began with fewer high-quality strains showed increases in partner quality because high-quality strains achieved higher relative fitness, whereas populations that began with many high-quality strains tended to decline in quality as those same strains had lower relative fitness (**Fig. 2**).

**Figure 2.**
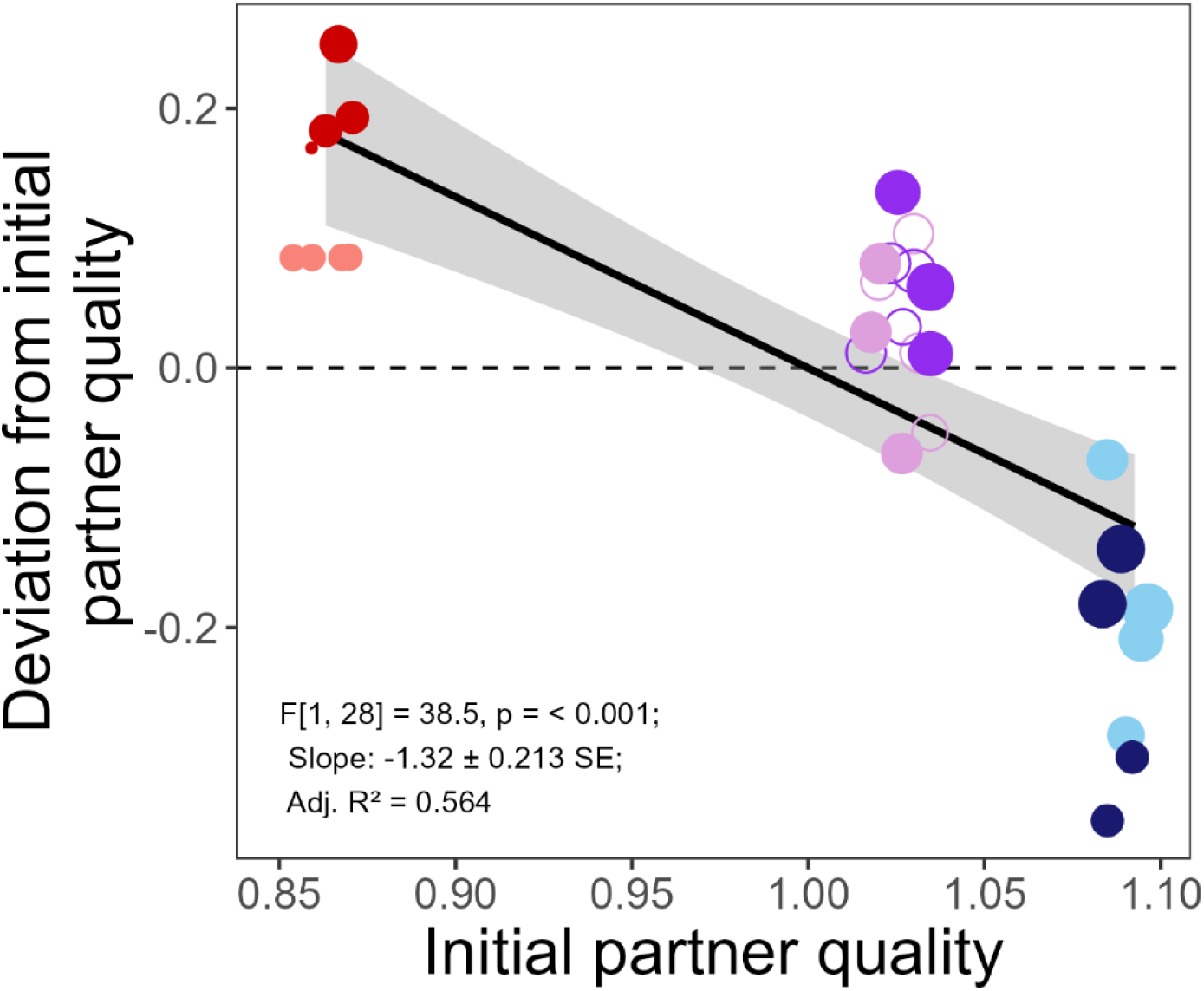
Negative frequency-dependent selection on rhizobium populations. Correlation between the mean partner quality values determined from the initial population strain compositions (x-axis) and the deviation from these values of the evolved population mean partner quality values (y-axis). Point size represents the percentage of strains recovered from each replicate population (i.e., soil slurry) at the end of the experiment, with colours representing the initial populations (Low-Quality = red; Medium-Quality = purple; High-Quality = blue) and shades the nitrogen environment (lighter = N-, darker = N+). Selective environments in which host plants were present or absent are represented by the closed and open points, respectively (note: host plant presence was manipulated only for the Medium-Quality populations). Population-level analyses used independently evolved replicate populations (soil slurries) as the unit of replication, with each point representing one replicate population (n = 30 total; n = 8–14 per treatment group).

### Evolutionary responses to selective environments

To determine how selection under contrasting environments shaped key mutualism traits across the three initial population types, we inoculated a new cohort of clovers (same accession used during experimental evolution) with slurries containing independently evolved replicate rhizobial populations harvested at the end of the evolution experiment. Plants were grown for ∼7 weeks under N-limitation (the same N-condition used during evolution), with each slurry used to inoculate multiple replicate plants (with plant-level measurements nested within replicate rhizobium populations). Nitrogen was not manipulated in this assay; thus, differences among populations reflect the effects of treatments experienced during the experimental evolution phase. We quantified shoot biomass as a measure of population-level partner quality, along with nodule number and per nodule weight as indicators of host investment. Unlike single-strain partner quality assays, these whole population assays capture the net functional outcome of evolutionary changes in strain frequencies, interactions, and population level traits.

Rhizobial evolutionary responses for nodulation traits depended on the initial population type (nodule number: χ^2^ = 4.74, *P* = 0.0936; per nodule weight: ^2^ = 8.62, *P* = 0.0135; **SI Appendix Table S3**). In contrast, for plant growth, the effects of population type depended on the nitrogen environment (Population Type × Nitrogen: shoot biomass: χ^2^ = 4.88, *P* = 0.0871; **Fig. 3**, **SI Appendix Table S3**). Although plants inoculated with rhizobia evolved under N+ environments tended to show 25.9% (95% CL: -51.6 to 13.4) less growth relative to those evolved under N- environments, consistent with the previous field study (37), this decline occurred only for the Medium-Quality populations and only when plant hosts were present during passaging (**Fig. 3**). The High- and Low-Quality populations showed the opposite trend: N+ environment-derived populations increased plant biomass by 17.2% (23.7 to 80) and 26.6% (-13.9 to 86.4), respectively, compared to those from N- environments. In the Medium-Quality populations evolved without hosts (soil-only treatments), the nitrogen environment populations experienced had little detectable effect on plant growth (-2.42%, -35.9 to 48.5; **Fig. 3**). Together, these patterns suggest that hosts can mitigate declines in partner quality under nitrogen-supplementation, but this host-mediated effect depends strongly on the initial population composition.

**Figure 3:**
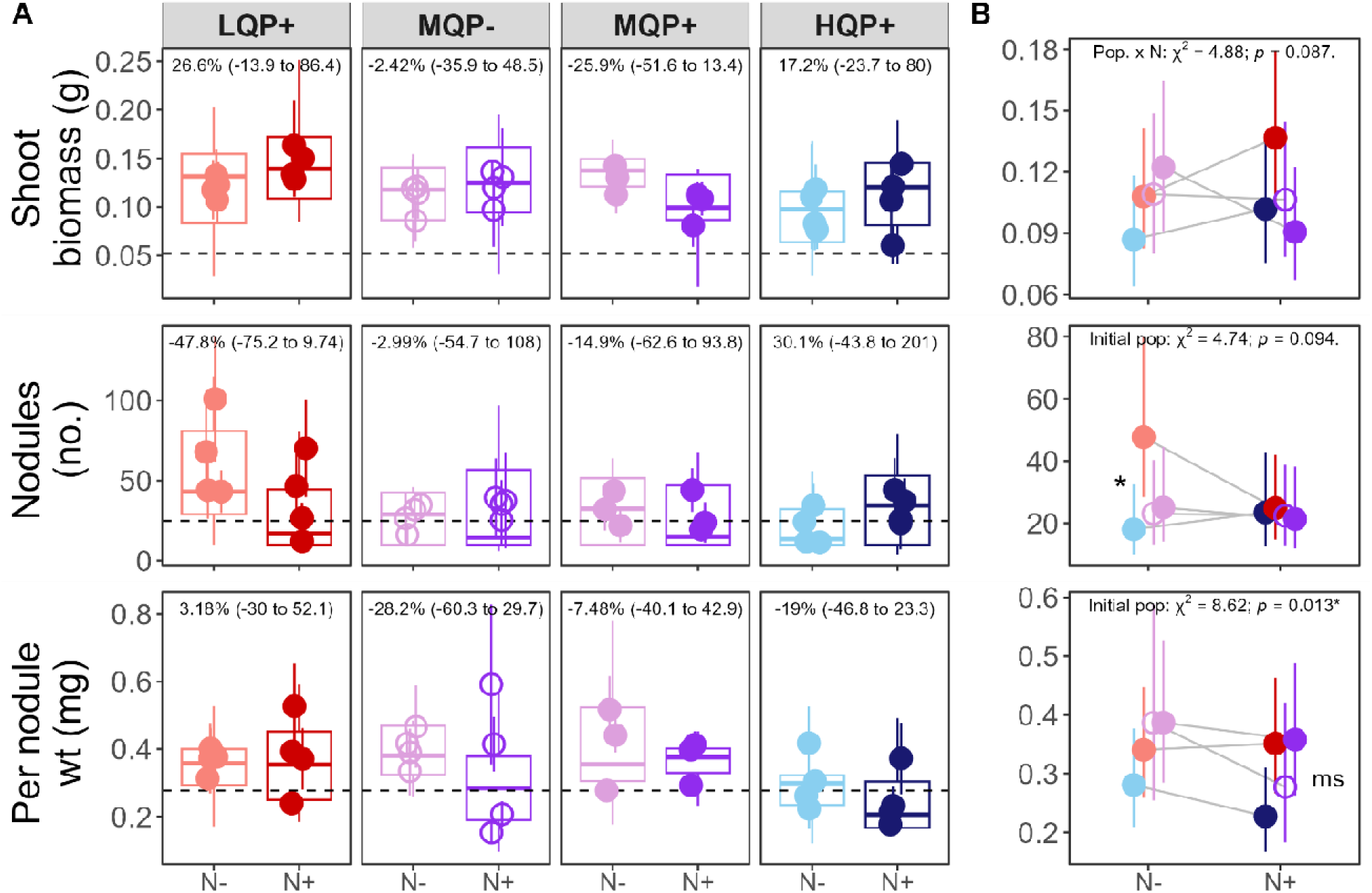
The evolutionary responses of rhizobia to selective environments depended on the initial population type. **A** Boxplots show plant-level responses from the soil slurry experiment, in which each soil slurry (representing an independently evolved replicate population) was used to inoculate multiple replicate plants randomized across treatments. Overlaid points and error bars show the mean +/- SE for each replicate population (i.e., averaged across plants inoculated with the same slurry; n = 3-4 populations per treatment). Horizontal dashed lines represent means for uninoculated controls. **B** Interaction plots show back-transformed estimated marginal means (EMMs) +/- 95% confidence limits (CLs) from linear mixed-effects models. Two models were fitted: one testing the effects of initial rhizobial population type and nitrogen treatment (using host-present treatments), and one testing the effects of host presence and nitrogen environment (using Medium-Quality populations only). Analyses included plant-level observations nested within replicate populations, with replicate population and tray position fitted as random effects (see **SI Appendix Table S6** for sample sizes corresponding to each analysis). Colours represent initial population type (“Low-Quality” in red, “High-Quality” in blue; and “Medium-Quality” in purple) while shades represent the nitrogen environment during experimental evolution (lighter = N-, darker = N+). Open and closed points indicate absence or presence of host plants during experimental evolution, respectively. Significant (*P* < 0.05) or marginally significant contrasts are indicated by an asterisk (*) or ms, respectively (see **SI Appendix Dataset S1**).

Nodulation traits showed the clearest evolutionary responses to nitrogen-supplementation, especially in the Low-Quality populations (**Fig. 3**). Relative to nitrogen-limited evolution, nitrogen-supplemented populations produced 47.8% fewer nodules (95% CL: -75.2 to 9.74), despite increasing plant growth. This pattern is consistent with evolved reductions in nodulation costs yielding greater net host benefit. Comparing among the initial population types within each N environment separately, the Low-Quality populations evolved to produce 164% more nodules (2.76 to 580) than the High-Quality population under N-limitation. Under the N+ environment, by contrast, the Low-Quality population evolved to produce 54.5% heavier nodules (-6.6 to 155) than the High-Quality population (**SI Appendix Dataset S1**). Both patterns indicate that the Low-Quality populations passaged under nitrogen-supplementation evolved traits that made them more beneficial to hosts, an effect not observed for the other population types.

### Impact of selection on standing genetic variation

To determine how selective environments and initial population types influenced the maintenance of genetic variation within replicate populations, we quantified the proportion of distinct strains recovered from nodules at the end of experimental evolution, with each replicate population treated as a single observation. We found that retention of genetic variation depended strongly on the initial population type (F [2,16] = 15.578, *P* < 0.001) as well as both selective environments (N: F [1,10] = 6.093, *P* = 0.033; host: F [1,10] = 7.411, *P* = 0.021; **Fig. 4**, **SI Appendix Table S4**). For the Medium- and Low-Quality populations, evolution under nitrogen-supplementation better preserved genetic diversity, with 22.0% (1.51 to 46.6) and 18.6% (-15.9 to 67.2) more strains retained on average compared to the same populations evolved under nitrogen-limitation (**Fig. 4**). Across N-environments, the Low-Quality populations consistently lost more genetic variation than the other population types, retaining 46.3% (-58.7 to -30.4) and 46.5% (-60.1 to -28.2) fewer strains compared to the Medium- and High-quality populations, respectively. In addition, in the Medium-Quality population, genetic diversity was 16% (2.73 to 31) higher when hosts were present during experimental evolution than when they were absent (**SI Appendix Dataset S2**). Together, these patterns suggest that while hosts and nitrogen environments were not direct agents of selection acting on strain frequencies, they nevertheless played important roles in shaping the amount of genetic variation that persisted through passaging by helping to preserve strain diversity, variation that is essential for enabling continued evolutionary responses in rhizobial populations.

**Figure 4:**
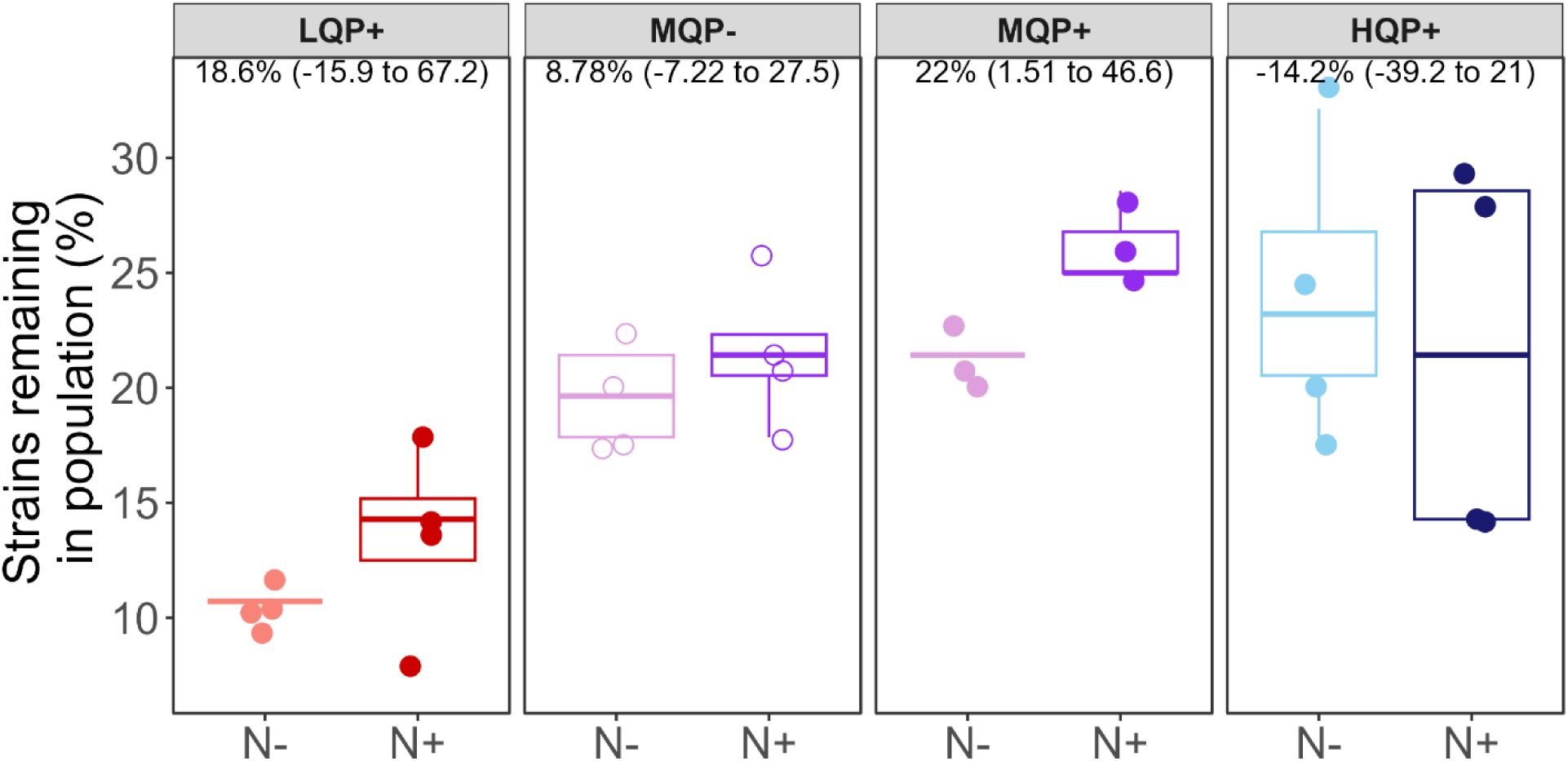
Nitrogen-supplementation and host presence preserves genetic variation during experimental evolution. Boxplots and points show the percent of strains retained at the end of experimental evolution for each replicate population (n = 30), estimated from nodule occupancy assays using 4 - 16 isolates sampled per replicate population. Colours denote initial population types (“Low-Quality” in red, “Medium-Quality” in purple, and “High-Quality” in blue) while shades represent the nitrogen environment during experimental evolution. Open versus closed points indicate whether host plants were absent or present during experimental evolution, respectively.

## Discussion

Despite predictions that selection for cooperation and cheating in mutualisms should erode genetic variation, natural populations consistently retain substantial diversity in partner quality. This unexpected genetic variation raises a fundamental question: what forces preserve this variation, and how do they shape the evolution and persistence of mutualisms? Here we use experimental evolution to reveal that selection on partner quality is not fixed but context-dependent, shifting with the genetic makeup of the initial population. This dynamic leads to negative frequency-dependent selection on partner quality, a mechanism that may play a crucial role in preserving genetic diversity in mutualist populations. While nitrogen alone was not a dominant selective force in our study, its contribution to maintaining diversity suggests it could buffer against host-driven selection for high-quality partners, thereby helping to sustain variation essential for mutualism stability.

### Negative frequency-dependent selection preserves mutualist variation

Negative frequency dependent selection (NFDS), whereby the per capita fitness of an individual genotype decreases as its relative abundance in a population increases (24, 43), is thought to be an important driver of the maintenance of both genetic and species diversity (43, 44). While well-characterized in host-pathogen and predator-prey systems (26, 45–47), empirical evidence for NFDS in mutualism is scant (48–50) and has only been explored by a small number of theoretical models (51, 52). By experimentally evolving multiple populations with differing initial genotype frequencies, in replicate, our work provides robust evidence for NFDS acting on partner quality.

Several theoretical models identify NFDS as a mechanism maintaining variation in mutualisms, particularly in interguild systems involving multiple host and partner genotypes. In such models, NFDS arises either because host genotypes compete for shared symbiont-derived resources (51) or because partner genotypes differ in the benefits they provide to different hosts, generating feedbacks that prevent any single partner type from dominating (52). For example, if mutualist hosts compete for shared symbiont-derived resources, then the fitness benefits of mutualism decline as cooperative host strategies become more common in the population (51). Alternatively, if the symbiont genotype most beneficial to host A is not the genotype most strongly favoured by host A, mismatched reciprocal preferences can generate negative feedbacks whereby common host genotypes preferentially enrich symbiont genotypes that disproportionally benefit competing host genotypes (52). In both cases, rare host or symbiont genotypes gain a relative fitness advantage, promoting the maintenance of variation through NFDS. These dynamics typically require a multispecies or multi-genotype framework, whereas our experimental design involved a single host genotype interacting with partner populations differing only in their starting frequencies of high- and low-quality strains.

Accordingly, our approach points to other mechanisms leading to NFDS, aligning with theory and empirical work exploring conditional host sanctions and allocation where host investment varies with the marginal fitness benefits of associating with high-quality partners (5, 25, 36). Our results suggest the following mechanism: when high-quality strains are rare, hosts gain large incremental benefits from each cooperative nodule and increase allocation or sanctioning effort. As high-quality strains become common and host nitrogen limitation is progressively alleviated, the marginal benefit of additional cooperative nodules declines, potentially relaxing host investment and allowing lower-quality strains to persist. This diminishing-returns dynamic generates NFDS and parallels a Volunteer’s Dilemma, in which once a minimal threshold of cooperators supplies the public good, additional cooperation yields smaller payoffs, enabling coexistence of cooperative and less-cooperative types (12).

### Environmental context as a driver of evolutionary capacity

Contrary to predictions that nitrogen-supplementation weakens mutualism by favouring low-quality rhizobia, we found little evidence that nitrogen-supplementation directly alters strain relative fitness, reinforcing the view that N is not always a strong direct selective agent on partner quality (53–55). On one hand, our finding of limited nitrogen effects on rhizobium evolution contrasts with prior work in this system documenting evolutionary declines in cooperation in response to more than two decades of field fertilization at KBS (37); however, the one condition where evolutionary declines were observed in our experiment (i.e., the Medium-Quality population with hosts present), was the condition most likely to resemble the original KBS soil community. A key difference between our design and the previous field study is that we isolated the effect of nitrogen-supplementation, removing covarying factors such as increased grass and forb growth, reduced light availability to legumes, and lower legume density (56, 57). These factors, rather than nitrogen alone, may drive declines in partner quality in natural settings (58).

Though it did not alter the direction of selection, nitrogen-supplementation nonetheless played a critical evolutionary role: it helped to maintain strain richness and preserve genetic diversity. This buffering effect explains two key observations. First, population-level partner quality remained robust across environments, likely because nitrogen-supplementation supported the persistence of high-quality strains within mixed populations, enabling hosts to associate with beneficial partners even when those populations evolved in the absence of hosts. This robustness in population-level partner quality aligns with evidence that population-level outcomes depend more on the presence of high-quality strains than on overall diversity (49, 53) (except see (59, 60)). Second, the Low-Quality populations evolved greater per-nodule benefits under nitrogen-supplementation. Why this shift occurred only under nitrogen-supplementation may reflect the role of fertilization in supporting larger rhizobial population sizes or reducing demographic loss, thereby preserving the genetic variation required for adaptive responses. This interpretation is consistent with prior findings that nitrogen-supplementation maintains higher strain richness and nucleotide diversity (37, 61). Alternatively, nitrogen may have indirectly influenced trait expression or competitive dynamics in ways that favoured more efficient nodulation. Regardless of mechanism, these results highlight that selection in mutualisms operates on multiple trait dimensions: population-level partner quality, per-nodule benefit, and host demand, and that environmental context modulates evolutionary potential by sustaining the diversity needed for these responses.

Our results also highlight that soil selection may shape strain frequency dynamics, although our ability to evaluate this is limited because soil-only (i.e., host absent) environments were included only for the Medium-Quality population. In this population, strain frequencies remained similar whether hosts were present or absent, suggesting that soil selection alone can produce evolutionary outcomes resembling those observed under host selection. Such similarity could arise if soil fitness and symbiotic fitness are genetically correlated, but evidence from other studies indicated these correlations may be weak or host-genotype dependent. For example, Burghardt and colleagues (20) found a weak positive correlation between soil and host fitness when using a choosy host genotype (A17), and no correlation when using a more permissive host (R108), suggesting genetic decoupling between soil and symbiotic fitness. In our experiment, correlated fitness effects could cause soil selection to dampen or reinforce host-mediated selection, but because we did not include soil-only treatments for the Low- and High-Quality populations, we cannot determine whether soil selection tends to produce NFDS, PFDS, or no consistent frequency dependence across population types. Overall, these results highlight that partner quality likely reflects a multidimensional suite of traits expressed in soil and symbiotic contexts, and that selection acting during the free-living stage may be an important, yet underappreciated, contributor to the maintenance or erosion of partner quality variation.

Together, these results help explain why declines in cooperation sometimes observed under field conditions do not always arise in controlled experiments: covarying ecological effects in natural settings, not nitrogen alone, likely drive losses of partner quality. More broadly, our findings show that environmental context modulates evolutionary potential rather than the target of selection. By maintaining standing genetic variation, nitrogen-supplementation can shape the tempo and magnitude of mutualism evolution, with ecological implications for the resilience of plant–microbe interactions under global change.

## Conclusions

Our results reveal that negative frequency-dependent selection allows high- and low-quality rhizobial strains to coexist, offering a robust explanation for the persistence of partner quality variation in mutualisms. Environmental conditions, such as nitrogen supplementation and host presence, did not alter selection but buffered genetic diversity, maintaining the raw material for adaptation. Together, these findings shift how we view mutualism stability: resilience arises not only from host control but from interactions between selection dynamics and symbiont diversity. This insight helps predict how plant–microbe partnerships will respond to global change drivers including nutrient deposition.

## Materials and Methods

### Study system

We used seeds of *Trifolium hybridum* (accession GRIN PI#419333), originally collected from Greece and previously used in single-strain inoculation experiments to estimate partner quality (37). *Trifolium* (i.e. clover) species typically form nodules with *Rhizobium leguminosarum* symbiovar *trifolii*, which carry the nod genes required for clover nodulation. These plant species produce indeterminate nodules, where rhizobia undergo terminal differentiation: the nitrogen-fixing bacteroid form becomes reproductively sterile, while undifferentiated cells remain viable when released from nodules. We used 56 rhizobia strains originally described as *R. leguminosarum* that had been isolated from field plots at the Kellogg Biological Station (KBS) Long-Term Ecological Research site (Michigan, USA) (37); half (n = 28) isolated from plots that had been annually supplemented with nitrogen fertilizer (ammonium nitrate) for over two decades (KBS Field N+), while the other half (n = 28) came from spatially paired plots that remained unfertilized for the same duration of time (KBS Field N-). However, long-read sequencing of these strains has since shown that multiple *Rhizobium* species are present (61).

### Experimental evolution study

Our experiment was designed to test two distinct hypotheses about the maintenance of partner quality variation in mutualism. To test our first hypothesis, that evolutionary outcomes depend on the starting frequency of high-quality partners and soil nitrogen availability, we established three population types (Low-Quality [LQ], Medium-Quality [MQ], and High-Quality [HQ]) and passaged them under two nitrogen environments (nitrogen-supplemented [N+] and nitrogen-free [N–]). While the three rhizobial population types showed distinct distributions of strain partner qualities prior to passaging (**SI Appendix Fig. S2A**), the growth benefits each provided when inoculated as a whole population did not necessarily differ (**SI Appendix Fig. S4,** growth cycle I). Each population type × nitrogen environment was propagated in 3-4 independently evolving replicate populations, yielding 22 populations in total (6 MQ [3 N-, 3 N+], 8 LQ [4 N-, 4 N+], 8 HQ [4 N-, 4 N+]; **SI Appendix Fig. S2B**). For the second hypothesis, we focused on the MQ population to disentangle the direct effects of nitrogen from host-mediated effects by additionally including replicate populations passaged without hosts (P-). Each host presence (P-, P+) × nitrogen environment (N-, N+) combination was propagated in 3-4 independently evolving replicate populations, yielding 14 MQ populations in total (8 P- [4 N-, 4 N+], 6 P+ [3 N-, 3 N+]; **SI Appendix Fig. S2B**). Because host presence was manipulated only within the MQ population, the overall design is not fully factorial across all population types. However, each treatment combination was represented by 3-4 independent replicate populations.

In total, the experiment comprised eight population type × nitrogen/host environment combinations and 30 independently evolving replicate populations. These replicate populations, each maintained in a single pot over four plant growth cycles (∼1 year; ∼400 rhizobial generations; **SI Appendix Fig. S2B**), constitute the primary unit of replication for all evolutionary analyses. Exceptions include genotypic selection analyses, which were conducted at the strain-level with observations nested within replicate populations, and plant-based assays, which were conducted at the plant level, with measurements nested within replicate populations (soil slurry experiment) or as repeated measures of pots across growth cycles (experimental evolution time series; **SI Appendix Table S6**). Replicate populations assigned to P+ treatments included surface-sterilized *T. hybridum* hosts. Each growth cycle consisted of a ∼6-week symbiosis phase followed by a ∼3-week free-living soil phase to allow for nodule senescence, after which roots were removed and (for P+ treatments) new seeds were sowed. No reinoculation occurred. At each cycle, shoot biomass was dried and weighed; nodules were destructively sampled from a subset of plants. See **SI Appendix Methods S1** for more details.

### Soil slurry experiment and isolate archiving

At the end of the final growth cycle, we prepared soil slurries from each replicate population (i.e., pot) by mixing homogenized soil with sterile saline and filtering through autoclaved cheesecloth to remove large soil particles. Slurries diluted out nutrient inputs while retaining rhizobia. Up to four replicate Conetainers™ (SC10R (62)) per slurry were filled with sterilized sand:turface mix and inoculated with 20 mL filtrate; controls received sterile saline. A single *T. hybridum* seed was sown into each pot (same accession used in experimental evolution). Pots were randomized across racks in the same chamber and conditions as the evolution experiment and plants grown for ∼6 weeks with weekly N-free Fahraeus fertilizer. We harvested plants destructively, recording shoot biomass and nodule number and mass. Approximately ten surface-sterilized nodules per plant were crushed and streaked onto TY agar, and a single purified colony per nodule was archived in 15% glycerol stocks stored at –80 °C.

### Short-read whole genome sequencing

From isolates archived after our experimental evolution study (derived isolates), we extracted DNA using a Bacterial Genomic DNA Isolation kit (BioVision: K309-100) following the manufacturer’s protocol. To confirm taxonomic identity and rule out contaminants, we amplified the full-length 16S rRNA gene of each isolate using universal primers 27F and 1492R under standard PCR conditions (63). PCR products (∼1500 bp) were verified by gel electrophoresis, purified with a GenElute™ PCR Clean-Up Kit (Sigma-Aldrich: NA1020), and submitted for Sanger sequencing at the Roy J. Carver Biotechnology Center (University of Illinois Urbana-Champaign, USA). We aligned forward and reverse reads using MEGA software (64) and identified taxa via NCBI’s Nucleotide BLAST (65). Of the ∼400 samples submitted for Sanger sequencing, 363 were confirmed as *Rhizobium* spp. We then submitted these verified *Rhizobium* DNA samples, representing 39–54 derived isolates per experimental evolution treatment or 4-16 isolates per replicate population, to the Roy J. Carver Biotechnology Center for Nextera XT library preparation and Illumina NovaSeq S1 sequencing (2 × 150 bp paired-end reads). Sequencing followed the center’s standard protocol.

### Long-read whole genome sequencing and assembly

Long-read sequencing of all 56 ancestral strains followed the protocol described in Vereau Gorbitz et al. (61). Briefly, isolates were grown on TY medium at 30 °C, and high-molecular-weight DNA was extracted using the PacBio Nanobind CBB kit. Library preparation and HiFi sequencing were performed at the W. M. Keck Center on a PacBio Sequel IIe system. Reads were processed with SMRT Link (v11.0), and genomes were assembled using the recommended Trycycler v0.5.5 workflow (66), including Filtlong filtering and assemblies generated by Flye, Hifiasm, and Raven. All assemblies were closed, and genome annotation was performed with NCBI PGAP v6.3.

### Determining the Most Probable Ancestor

Illumina reads from 363 derived isolates were quality-filtered, assembled, and assessed using standard bacterial genomics workflows (full commands and software versions in **SI Appendix Methods S2**). Genome quality and completeness were high for nearly all isolates, except for a few contaminated or taxonomically ambiguous genomes (**SI Appendix Dataset S3**). Species identification was performed using NCBI’s PGAP Taxonomy pipeline, which classified 359 of 363 genomes as *Rhizobium leguminosarum*. To identify the most probable ancestor (MPA) for each derived isolate, we calculated pairwise average nucleotide identity (ANI) between each isolate and all 56 ancestral strains. The ancestral strain with the highest ANI was designated as the MPA. Most isolates showed clear MPA assignments, with the best ANI match typically exceeding the second-best by a large margin (**SI Appendix Fig. S3**). In the few cases where two candidates had nearly identical ANI values, we resolved ties using the known population of origin (LQ, MQ, HQ). Only a single isolate showed evidence of cross-contamination. Overall, ANI-based assignments were highly resolved, allowing confident identification of 26 distinct MPAs across the 363 isolates.

### Statistical analyses

We analyzed selection and evolutionary responses using complementary approaches that differ in their unit of replication. We distinguish between (i) strain-level analyses, (ii) population-level analyses, and (iii) plant-level assays (**SI Appendix Table S6**).

*(i) Strain-level analyses* quantify selection within replicate populations, with strain-level observations nested within populations. For these analyses, the number of observations reflects the number of strains (28 per population) multiplied by the number of replicate populations within each dataset (e.g., 28 x 22 = 616 observations for the full host-present dataset; see **SI Appendix Table S1**). To test how selection acted on rhizobial strains under different selective environments (**SI Appendix Methods S3i**), we estimated strain-level fitness from changes in strain frequency and paired these values with previously measured partner-quality (PQ) for each strain (37). We fitted global models to test whether selection differed across experimental factors (e.g., trait x population type or trait x environment interactions). When selection varied among factors (e.g., population type), we measured within-group non-linear quadratic selection coefficients (γ) from models including linear and non-linear trait terms and estimated directional selection differentials (*S*) representing total selection on the trait, from separate univariate linear models. Models initially included replicate population as a random effect; however, this term explained negligible variance and was therefore omitted from final models.
*(ii) Population-level analyses* use independently evolving replicate populations as the unit of replication (n = 8-14 per group: 30 total). To evaluate how PQ distributions shifted during experimental evolution, we compared initial and final PQ values within replicate populations using paired t-tests. We tested for negative frequency-dependent selection by regressing each replicate population’s change in mean PQ (evolved − initial) against its initial mean PQ. We further compared observed changes to null expectations generated by resampling strain-level partner quality values within replicate populations, matched to the observed sampling effort. Finally, we tested how genetic diversity (the proportion of strains recovered per replicate population) depended on population type and selective environment using linear models (**SI Appendix Methods S3iii)**.
*(iii) Plant-based assays* were conducted at the level of individual plants, with observations nested within replicate populations (soil slurry experiment) or within pots measured repeatedly across growth cycles (experimental evolution time series; reported in **SI Appendix Results**). To assess how selective environments affected mutualism traits measured in the soil slurry experiment (**SI Appendix Methods S3ii**), we fitted linear mixed-effects models with fixed effects corresponding to the dataset analyzed (see **SI Appendix Table S6**), including nitrogen environment and either population type (P+ dataset) or host presence (MQ dataset), with replicate population (soil slurry) and tray position as random effects. Significant effects were followed by post hoc contrasts with multiple-testing correction.

### Disclosure of Delegation to Generative AI

The authors declare the use of generative AI in the research and writing process. According to the GAIDeT taxonomy (67), the following tasks were delegated to GAI tools under full human supervision: Code optimization, and Proofreading and editing. The GAI tool used was: Co-pilot GPT 5.2. Responsibility for the final manuscript lies entirely with the authors. GAI tools are not listed as authors and do not bear responsibility for the final outcomes. Declaration submitted by: RTD.

## Acknowledgements and funding sources

We thank members of the Heath, diCenzo, and Doyle Labs, as well as McMaster Biology’s Bioinformatics, Evolution, Anthropology, and Population Genetics Seminar attendees for their feedback on early drafts of this manuscript. We thank Laura Goralka for assistance in the greenhouse. We acknowledge the Roy J. Carver Biotechnology Center for their valuable technical support and guidance, with special thanks to Alvaro G. Hernandez and Chris L. Wright. This work was supported by funding from the following sources: Carl R. Woese Institute for Genomic Biology, Arnold O. Beckman Research Award (CRB-RB22083: University of Illinois, Urbana-Champaign), the National Science Foundation (IOS-1645875, DBI-2022049, PGRP-1856744), Natural Sciences and Engineering Research Council of Canada Discovery Grants Program (RGPIN-2023-05144, DGECR-2023-00046), Canadian Foundation for Innovation John R. Evans Leaders Fund (42856), and McMaster University.

## Author contributions

Conceptualization (RTD, KDH, JAL), Data curation (RTD, XS, CG, OO, KG), Formal analysis (RTD, XS), Funding Acquisition (RTD, KDH), Investigation (RTD, MB, KG, OO, CG, EP, XS), Methodology (RTD, XS), Project Admin (RTD), Resources (RTD, KDH, DVG), Software (RTD, XS), Supervision (RTD, KDH), Validation (RTD, XS, EP), Visualization (RTD, XS), Writing (original draft: RTD, KDH; review and editing: RTD, KDH, JAL).

## Competing Interest Statement

the authors declare no competing interests

## Notes

### Competing Interest Statement

The authors have declared no competing interest.

### Summary of Updates

More explicit descriptions of experimental design, replication structure, and statistical analyses.

