## Supplementary material for "Negative frequency-dependent selection maintains partner quality variation in a keystone nutritional mutualism": SI Appendix

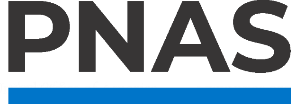

**Supporting Information for**

### Negative frequency-dependent selection maintains partner quality variation in a keystone nutritional mutualism

**Authors:**

Rebecca T. Doyle^1,2*^, Xingyuan Su^1^, Christina Gallick^2,3^, Meghan Blaszynski^2,3^, Emily Perry^1^, Karla Griesbaum^3^, Omolabake O. Oyetayo^3^, David Vereau Gorbitz^2,3^, Jennifer A. Lau^5^, Katy D. Heath^2,3,4^

**This PDF file includes:**

Methods S1-S3

Results S1

Tables S1 to S6

Figures S1 to S4

Legends for Datasets S1 to S4

SI References

**Other supporting materials for this manuscript include the following:**

Datasets S1 to S4

#### Supporting Methods

###### SI Appendix Method S1: Experimental evolution study

###### i. Rhizobial populations and starting composition construction

The three initial populations differed in the starting frequencies of high-quality strains, resulting in three distributions of partner‑quality phenotypes:

1. High‑Quality (HQ): 28 strains from KBS N– field plots (*μ* = 0.500, σ = 0.136);
2. Medium‑Quality (MQ): 14 N– + 14 N+ strains (*μ* = 0.469, σ = 0.166); and
3. Low‑Quality (LQ): 28 strains from KBS N+ plots (*μ* = 0.395, σ = 0.162).

Strains were grown individually in TY broth (29°C, 200 rpm) (1), normalized to OD600 ≈ 0.1, and pooled in equal volumes before inoculating 35 mL into 1‑L pots containing autoclaved low‑nutrient soil (1:1 Berger:turface (2, 3)), yielding ~3.5 × 10⁷ cells/g soil.

###### ii. Passaging design, replication structure, and growth‑chamber layout

Each of the HQ, MQ, and LQ populations evolved in N− and N+ environments, with MQ additionally split into P+ and P– treatments (**SI Appendix Fig. S2B**). Replication: four independently evolving replicate populations per treatment, except MQ P+ (three replicate populations). Pots (N = 30) were distributed across pre‑bleached trays (six trays for HQ and LQ, eight for MQ), with each tray containing: 3 pots assigned to each of the two nitrogen treatments (N+ and N–) plus one uninoculated control, for a total of seven pots per tray. MQ trays additionally included two P– pots (one N+ and one N–). Trays were randomized weekly within a controlled growth chamber (16:8 light:dark; 21:18°C; 60% humidity; 300 μmol/m²/s PAR). Pot bottoms were enclosed in autoclave bags to prevent cross‑contamination. For P+ treatments, three surface‑sterilized *T. hybridum* seeds were sown.

###### iii. Nutrient conditions

All pots received N‑free Fahraeus solution (4) for macro/micronutrients. N+ treatments followed the KBS fertilization regime (5) scaled to pot size (0.66 g NH₄NO₃ per pot per cycle). Within each cycle, nitrogen concentration increased gradually to avoid seedling shock (50 ppm to 850 ppm over 5 weeks).

###### iv. Passaging procedure

Each cycle proceeded as:

1. Symbiosis phase (6 weeks): Pots inoculated with mixed rhizobia.
2. Induced senescence: Shoots cut to stimulate nodule breakdown.
3. Free‑living soil phase (~3 weeks): Rhizobia released from nodules into soil.
4. Next cycle: Roots removed, soil rehydrated; seeds replanted (P+ only).

No reinoculation occurred, ensuring evolution proceeded from existing soil populations with negligible gene flow.

###### v. Phenotyping during evolution (time series)

At each growth cycle, shoots were dried (60°C, 3 days) and weighed, and a subset of five haphazardly chosen plants per treatment had roots harvested for nodule counts and nodule biomass.

###### SI Appendix Method S2: Determining the most probable ancestor (MPA)

###### i. Read processing and genome assembly

Illumina paired‑end reads for 363 derived isolates were processed on the McMaster Biology Department’s high‑performance server. All Linux commands, R scripts, and outputs are available on GitHub (to be made public upon publication; accessible to reviewers upon request), and raw reads have been deposited in NCBI’s Sequence Read Archive (BioProject id: PRJNA310138). Read quality was assessed with FastQC v0.12.1 (6), and summary reports were generated with MultiQC v1.28 (7). Reads were trimmed and filtered with fastp v0.23.4 (8). Genomes were assembled with SPAdes v3.15.5 (9) in “isolate” mode using paired, merged, and unpaired reads. Assembly quality metrics (scaffold count, genome size, N50, GC content) were computed using QUAST v5.2.0 (10), and completeness/contamination were evaluated with CheckM v1.2.3 (11) using lineage‑specific marker sets. Species identification was performed using NCBI PGAP Taxonomy v2025‑05‑06 (build 7983), which compares query genomes to type strains via pairwise average nucleotide identity (ANI) and coverage. Mean assembly statistics across isolates were: 134 scaffolds, genome size ~7.5 Mb, N50 ~514 kb, and GC content ~60.6% (**SI Appendix Dataset S3**). Nearly all genomes were >99% complete with <5% contamination, except for two genomes with ≥100% contamination and mixed GC content. PGAP classified 359 of 363 isolates as *R. leguminosarum* (358 high confidence; one low confidence). We also detected one contaminated genome (still assigned to *R. leguminosarum*), two isolates with inconclusive species assignments (*Rhizobium viscosum* and *Paenibacillus allorhizosphaerae*), and one isolate assigned to *Rhizobium johnstonii*.

###### ii. ANI‑Based Most Probable Ancestor Assignment

Pairwise ANI between each derived isolate and all 56 ancestral strains was computed using FastANI v1.32 (12). The ancestral strain with the highest ANI value was designated the most probable ancestor (MPA). The mean ANI of the top hit was 99.9976 (SD = 0.0013), far exceeding the second‑best hit (mean = 99.0821, SD = 0.5202; **SI Appendix Fig. S3**). In four cases where two candidate MPAs differed by <0.001 ANI, ties were resolved using known population type (LQ, MQ, HQ). In one instance, a top ANI hit originating from a mismatched population was interpreted as cross‑contamination. In total, the 362 derived isolates were assigned to 25 MPAs.

###### SI Appendix Method S3. Statistical analyses

All analyses and figures were conducted or created in R (13), using the following packages: tidyverse v2.0.0 (14), knitr v1.50 (15), kableExtra v1.4.0 (16), lme4 v1.1-37 (17), lmerTest v3.1-3 (18), car v3.1-3 (19), emmeans v1.11.2-8 (20), cowplot v1.2.0 (21), and viridis v0.6.5 (22).

###### i. How does selection act on rhizobium populations that evolved under distinct selective environments?

**Fitness calculation and trait standardization:** We calculated strain frequencies within each replicate soil-slurry population by dividing the number of isolates assigned to each strain by the total number of isolates recovered from that replicate population (4-16 isolates sampled per replicate population; N = 363 total isolates). Strain fitness was calculated as the log₂ fold change between its final nodule frequency and its initial inoculum frequency (~1/28). Strains not recovered from final populations were assigned a fitness value of 0, representing a reduction below the lowest detectable value. Fitness values were then relativized by dividing each strain’s fitness by the mean fitness of all strains within the same selective environment. Partner quality (PQ) estimates were taken from Weese et al. (5) and standardized across all 56 ancestral strains (mean = 0, SD = 1).

**Selection Analyses**: We performed genotypic selection analyses following Lande & Arnold (23) and Stinchcombe et al. (24). To test whether selection differed across experimental conditions, we first fitted global models including linear and quadratic terms and their interactions with experimental factors, with strain-level observations nested within replicate populations. These models took two forms: (i) fitness ~ (trait + trait^2^) x population type + (trait + trait^2^) x nitrogen environment, using data from the host‑present (P+) treatments; or (ii) fitness ~ (trait + trait^2^) x host environment + (trait + trait^2^) x nitrogen environment, using only strains from the Medium-Quality populations. These models were used to assess whether linear or non-linear selection varied across population types or selective environments, based on the significance of interaction terms. When no significant interactions were detected (e.g., for nitrogen environment), selection was interpreted as consistent across those conditions based on the global model.

When significant trait x factor interactions were detected (e.g., for population type), we estimated selection gradients within each level of that factor by fitting separate models of the form: fitness ~ trait + trait^2^. Quadratic (*γ*) selection gradients, which indicate non-linear selection, were obtained from these within-group models, with quadratic coefficients doubled after fitting to obtain canonical quadratic selection gradients (24). Linear selection differentials (*S*), representing total selection on a trait including indirect effects arising from correlated traits, were estimated separately from univariate models of the form fitness ~ trait. All reported selection gradients (*γ*) and differentials (*S*) reflect fitness differences accumulated over many rhizobial generations rather than single‑generation measures.

**Testing evolutionary change in PQ**: For each replicate population, we compared initial and final PQ values using paired t‑tests, with analyses performed separately for each population type (LQ, MQ, HQ). To test whether recovered strains were unusually high or low in PQ, we generated a null distribution for each replicate by randomly sampling (with replacement) the same number of strains for which occupancy was determined, calculating the median PQ of the sampled strains, and repeating this sampling 1,000 times. Observed median PQ values were compared to null distributions using empirical quantiles (2.5% and 97.5%) of the resampled distributions.

**Negative Frequency‑Dependent Selection on PQ**: We plotted each replicate population’s initial mean PQ against the deviation between evolved and initial PQ values. Evolved PQ values were expressed as proportional change relative to initial PQ: (evolved – initial) / initial. Populations with higher initial mean PQ (reflecting greater representation of high-quality strains) fall to the right on the x‑axis. Populations above zero on the y‑axis had increases in PQ; those below zero had decreases. A linear regression tested whether deviations in PQ were significantly associated with initial PQ, with a negative slope indicating negative frequency‑dependent selection.

###### ii. How does experimental evolution under distinct selective environments impact key mutualism traits?

We analyzed evolutionary responses from the soil slurry experiment using linear mixed‑effects models (LMMs). Because multiple plants were inoculated with each independently evolved population (soil slurry), plant-level observations were modeled with replicate population included as a random effect, ensuring that inference reflects the level of independent evolutionary replication. Two parallel model sets mirrored those used in the selection analyses: (i) population type × nitrogen environment experienced during experimental evolution (N−, N+), using host‑present (P+) replicate populations; and (ii) host presence × nitrogen environment, using Medium-Quality populations only. All traits were log‑transformed to meet model assumptions. Fixed effects included population type, nitrogen environment, and/or host presence depending on the dataset analyzed. Random effects included replicate population (soil slurry) and tray position in the growth chamber (20 trays total). Significance was assessed using Type III sums of squares (*car*::Anova). Post‑hoc comparisons were conducted using emmeans, with pairwise contrasts between nitrogen environments within each population type, or among initial population types within each selective environment. Effect sizes were calculated as log_2_ response ratios, and p‑values were adjusted for multiple comparisons using Dunnett’s (25) or Tukey’s method, for nitrogen environment or population type comparisons, respectively.

###### iii. What impact does experimental evolution under distinct selective environments have on standing genetic variation?

To assess how selective environments influenced the retention of genetic diversity within replicate populations, we used linear models testing whether the percentage of strains recovered from each final population depended on initial population type, nitrogen environment, and/or host presence. Analyses were subset to P+ treatments to test population × nitrogen interactions, and to MQ populations to test host × nitrogen interactions, paralleling the fixed‑effect structure of the LMM analyses above.

###### iv. How did plant growth change over the course of experimental evolution?

We analyzed shoot biomass over time using a LMM implemented with lme4, which tested the effects of inoculation treatment (four levels: LQ, MQ, HQ, uninoculated controls), contemporary nitrogen supplementation (N-, N+), plant growth cycle (four levels), and all two- and three-way interactions. Tray (n = 20) was included as a random effect. We log-transformed biomass to improve normality and tested the significance of fixed effects terms using type III sums of squares (*car*::Anova). We then computed estimated marginal means using emmeans and applied Dunnett’s contrasts to compare inoculated treatments against uninoculated controls (N- treatment only), as well as pairwise contrasts among population types within each nitrogen treatment and growth cycle, adjusting p-values for multiple comparisons.

#### Supporting Results

###### SI Appendix Result S1: Plant growth responses during experimental evolution.

We assessed the impact of inoculation treatment, contemporary nitrogen supplementation, and growth cycle, on plant growth, measured as dried above-ground (shoot) biomass. Our model showed significant main effects of all three terms, in addition to a significant interaction between N-supplementation and growth cycle (**SI Appendix Table S5**). The most pronounced effect of inoculation treatment on plant growth occurred in the final growth cycle in the N+ treatment, where plants inoculated with Low‑Quality populations in the first growth cycle grew 86% (95% CL = 0.8 to 242%) larger than those initially inoculated with High-Quality populations (**SI Appendix Fig S4**, **SI Appendix Dataset S4**). Across all growth cycles, uninoculated controls remained consistently smaller (range: 44 – 79%) than any inoculated plants (**SI Appendix Dataset S4**). Although nodules were found on controls in each cycle, these nodules did not improve plant growth, suggesting ineffective nitrogen fixation. Importantly, genetic screening showed virtually no cross‑contamination among populations (see Main Text **Materials and Methods**, *Determining the most probable ancestor*), indicating that nodules on control plants likely arose from within-population type contamination (i.e., among replicate populations derived from the same initial strain pool). Gene flow, if present, was therefore limited to within population types.

#### Supporting Tables

###### SI Appendix Table S1: Global analyses revealed stabilizing selection on partner quality across populations, with selection differing among initial population types.

Results of regression models testing whether directional (*β*) and quadratic (*γ*) selection on partner quality (PQ) varied across experimental factors. Analyses were conducted on two data subsets: (i) host-present (P+) populations to test effects of initial population type and nitrogen environment (N-, N+), and (ii) Medium-Quality populations to test effects of host presence (P-, P+) and nitrogen environment. We report Type III sums of squares (T3SS), degrees of freedom (Df), F‑statistics, and P‑values from ANOVA.

| **Model 1: Does selection operate differently depending on initial population type or N environment? Limit data to host-present (P+) data** | | | | | |
| --- | --- | --- | --- | --- | --- |
| **Model term** | **T3SS** | **Df** | **F** | ***P* (F)** | **Est. [+/- SE]** |
| Partner quality  (*β*) - PQ | 2.886 | 1 | 0.519 | 0.472 | 0.154 [0.214] |
| Partner quality  (*γ*) – PQ^2^ | **45.055** | **1** | **8.1** | **0.005** | **-0.936 [0.329]** |
| Population Type (pop) | 6.942 | 2 | 0.624 | 0.536 | NA |
| N environment (N) | 0.265 | 1 | 0.048 | 0.827 | NA |
| PQ x pop | **62.84** | **2** | **5.65** | **0.004** | NA |
| PQ^2^ x pop | 12.786 | 2 | 1.15 | 0.318 | NA |
| PQ x N | 0.47 | 1 | 0.084 | 0.771 | NA |
| PQ^2^ x N | 0.625 | 1 | 0.112 | 0.738 | NA |
| Residuals | 3359.232 | 604 | NA | NA | NA |
| **Model 2: Does selection operate differently depending on host and N environments? Limit to Medium-Quality population data** | | | | | |
| **Model term** | **T3SS** | **Df** | **F** | ***P* (F)** | **Est. [+/- SE]** |
| Partner quality  (*β*) - PQ | 0.084 | 1 | 0.020 | 0.887 | 0.0257 [0.180] |
| **Partner quality**  **(*γ*) – PQ^2^** | **40.383** | **1** | **9.780** | **0.002** | **-0.943 [0.302]** |
| Host environment (Host) | 0.171 | 1 | 0.041 | 0.839 | NA |
| N environment (N) | 0.016 | 1 | 0.004 | 0.951 | NA |
| PQ x Host | 0.321 | 1 | 0.078 | 0.780 | NA |
| PQ^2^ x Host | 0.326 | 1 | 0.079 | 0.779 | NA |
| PQ x N | 0.081 | 1 | 0.020 | 0.888 | NA |
| PQ^2^ x N | 0.027 | 1 | 0.006 | 0.936 | NA |
| Residuals | 1581.587 | 383 | NA | NA | NA |

###### SI Appendix Table S2: The direction and strength of natural selection acting on rhizobium populations during experimental evolution depended on the initial population type.

Total linear selection differentials (*S*), and non-linear quadratic (*γ*) selection gradients for partner quality (PQ), estimated using regression models fitted within each population type (Low-, Medium-, and High-Quality populations) following global tests for heterogeneity in selection. Estimates are shown with associated confidence intervals. Quadratic regression coefficients were doubled after model fitting to obtain canonical quadratic selection gradients (*γ*).

|  | ***S*** | | ***γ*** | |
| --- | --- | --- | --- | --- |
| **Population** | **ANOVA** | **Est. [+/- SE]** | **ANOVA** | **Est. [+/- SE]** |
| Low Quality | **F [1,222] = 6.61,**  **p = 0.011** | **0.508 [0.197]** | **F [1,221] = 6.44,**  **p = 0.0119** | **-0.878 [0.346]** |
| Medium Quality | **F [1,390] = 6.7,**  **p = 0.01** | **0.266 [0.103]** | **F [1,389] = 24,**  **p < 0.001** | **-0.884 [0.181]** |
| High Quality | **F [1,222] = 19,**  **p < 0.001** | **-0.675 [0.155]** | F [1,221] = 0.352,  p = 0.554 | -0.189 [0.318] |

###### SI Appendix Table S3: Linear mixed effects models (LMMs) testing evolutionary responses to selective environments.

Results from two sets of linear mixed-effects models (LMMs) evaluating how evolutionary responses of rhizobial populations (measured in the soil‑slurry experiment) depended on selective environment. The first model set tested the main and interactive effects of population type (Low-, Medium-, High-Quality) and nitrogen environment (N–, N+) experienced during experimental evolution using data from replicate populations evolved with hosts present (P+). The second model set tested the main and interactive effects of nitrogen environment (N–, N+) and host presence during evolution (P–, P+), using data from the Medium‑Quality population only. All models included random effects of (i) replicate population (soil slurry) and (ii) tray position in the growth chamber (20 trays total). Traits were log‑transformed to meet ANOVA assumptions, and the significance of fixed effects was assessed using Type III sums of squares (*car*::Anova). The table reports χ² statistics with corresponding p‑values in parentheses. Contrasts from models reported in **SI Appendix Dataset S1**.

| **Model 1: ANOVA results for LMMs testing the phenotypic evolutionary responses to initial population type, N environment, and their interaction. Limited to host-present (P+) data** | | | |
| --- | --- | --- | --- |
| **Trait** | **Initial population (Pop.)** | **N environment (N)** | **Pop x N** |
| Shoot wt | 3.83 (0.147) | 0.0842 (0.772) | *4.88 (0.0871)* |
| Nodule no. | *4.74 (0.0936)* | 0.754 (0.385) | 3.23 (0.198) |
| Nodule wt | **8.62 (0.0135)** | 0.608 (0.435) | 0.82 (0.664) |
| **Model 2: ANOVA results for LMMs testing the phenotypic evolutionary responses to host presence, N environment, and their interaction. Limited to Medium-Quality population data** | | | |
| **Trait** | **Host presence (Host)** | **N environment (N)** | **Host x N** |
| Shoot wt | 0.0196 (0.889) | 1.65 (0.199) | 1.31 (0.252) |
| Nodule no. | 0.0078 (0.93) | 0.0383 (0.845) | 0.00535 (0.942) |
| Nodule wt | 0.417 (0.518) | 1.02 (0.312) | 0.444 (0.505) |

###### SI Appendix Table S4: Linear models testing the effects of selective environments on post-evolution genetic diversity.

Results from two sets of linear models (LMs) that assessed whether genetic diversity after passaging, measured as the proportion of strains recovered per replicate population, depended on selective environment. The first model set evaluated the main and interactive effects of rhizobial population type (Low-, Medium-, High-Quality populations) and nitrogen environment (N–, N+) using data from replicate populations evolved with hosts present (P+). The second model set evaluated the main and interactive effects of nitrogen environment (N–, N+) and host presence during evolution (P–, P+), using data from the Medium‑Quality population only. For each model, we report Type II sums of squares (T2SS), degrees of freedom (Df), F‑statistics, and P‑values from ANOVA. Contrasts from models reported in **SI Appendix Dataset S2**.

| **Model 1: ANOVA results for LMs testing the impacts of initial population type, N-environment, and their interaction on standing genetic variation remaining at the end of experimental evolution. Limited to host-present (P+) data** | | | | |
| --- | --- | --- | --- | --- |
| **Model term** | **T2SS** | **Df** | **F** | ***P* (F)** |
| Initial pop. | **2.245** | **2** | **15.578** | **< 0.001** |
| N-environment | 0.020 | 1 | 0.278 | 0.605 |
| Pop x N | 0.144 | 2 | 0.999 | 0.390 |
| Residuals | 1.153 | 16 | NA | NA |
| **Model 2: ANOVA results for LMs testing the impacts of host presence, N-environment, and their interaction on standing genetic variation remaining at the end of experimental evolution. Limited to Medium-Quality population data** | | | | |
| **Model term** | **T2SS** | **Df** | **F** | ***P* (F)** |
| Host presence | **0.076** | **1** | **7.411** | **0.021** |
| N-environment | **0.062** | **1** | **6.093** | **0.033** |
| Host x N | 0.011 | 1 | 1.103 | 0.318 |
| Residuals | 0.102 | 10 | NA | NA |

###### SI Appendix Table S5: Linear mixed‑effects model (LMM) testing the effects of initial population type, nitrogen environment, and growth cycle on plant shoot biomass.

Results from a LMM evaluating how inoculation treatment (Low-, Medium-, and High-Quality population types, plus uninoculated controls), contemporary nitrogen supplementation (N–, N+), and plant growth cycle (four cycles total), along with all two‑ and three‑way interactions, influenced shoot biomass over the course of experimental evolution. Biomass was log‑transformed to improve normality, and the model included pot (replicate population, n = 8-20 per condition and growth cycle) and tray (n = 20) as random effects. Fixed‑effects significance was assessed using Type III sums of squares (car::Anova). The table reports degrees of freedom (Df), χ² statistics, and corresponding P‑values for each model term. Estimated marginal means and Dunnett‑adjusted contrasts used for post‑hoc comparisons are summarized in **SI Appendix Fig. S4** and **SI Appendix Dataset S4**, respectively.

| **Term** | **Df** | **χ^2^** | ***P* (χ^2^)** |
| --- | --- | --- | --- |
| **N environment (N)** | **1** | **47.00** | **< 0.001** |
| **Population (Pop)** | **3** | **24.20** | **< 0.001** |
| **Growth cycle (GC)** | **3** | **8.73** | **0.033** |
| N x Pop | 2 | 2.08 | 0.354 |
| **N x GC** | **3** | **21.20** | **< 0.001** |
| Pop x GC | 9 | 12.00 | 0.216 |
| N x Pop x GC | 6 | 9.17 | 0.164 |

###### SI Appendix Table S6: Summary of analytical approaches, units of replication, and sample sizes used across all models. For strain-level analyses (orange), observations reflect the number of strains per replicate population (n = 28) multiplied by the number of replicate populations included in each dataset. For population-level analyses (blue), observations correspond to the number of replicate populations. For plant-based assays, observations reflect plant-level measurements nested within replicate populations in the soil slurry experiment (light green) or repeated measurements of replicate populations (i.e., pots) across growth cycles in the evolution experiment (dark green).

| **Analysis** | **Dataset** | **Unit of replication** | **Design (factors)** | **Replication structure** | **N (obs.)** | **Reference** |
| --- | --- | --- | --- | --- | --- | --- |
| 1) Global selection model (LM) | P+ | Strain (nested in rep. pop.) | 3 pop. types × 2 N envs. | 22 rep. pops. × 28 strains | 616 | SI Appendix Table S1 |
| 2) Global selection model (LM) | MQ | Strain (nested in rep. pop.) | 2 N envs.× 2 host envs. | 14 rep. pops. × 28 strains | 392 | SI Appendix Table S1 |
| 3) Within population type selection models (LM) | LQ/HQ | Strain (nested in rep. pop.) | Within pop. type (pooled across envs.) | 8 rep. pops. × 28 strains | 224 | Fig. 1A, SI Appendix Table S2 |
| 4) Within population type selection models (LM) | MQ | Strain (nested in rep. pop.) | Within pop. type (pooled across envs.) | 14 rep. pops. × 28 strains | 392 | Fig. 1A, SI Appendix Table S2 |
| 5) PQ shifts  (t-tests) | LQ | Rep. pop. | Change in PQ (evolved − initial) | 8 rep. pops. | 8 | Fig. 1B |
| 6) PQ shifts  (t-tests) | HQ | Rep. pop. | Change in PQ (evolved − initial) | 8 rep. pops. | 8 | Fig. 1B |
| 7) PQ shifts  (t-tests) | MQ | Rep. pop. | Change in PQ (evolved − initial) | 14 rep. pops. | 14 | Fig. 1B |
| 8) NFDS (linear regression) | All | Rep. pop. | Initial PQ vs proportional change in PQ [(evolved - initial) / initial] | 30 rep. pops. | 30 | Fig. 2 |
| 9) Permutation tests | All | Rep. pop. | Observed PQ vs empirical null dist. | 30 rep. pops. | 30 | SI Appendix Fig. S1 |
| 10) Evolutionary responses (LMMs) | P+ | Plant (nested in rep. pop.) | 3 pop. types × 2 N envs. | 22 rep. pops. × 1–4 plants | 75 | Fig. 3, SI Appendix Table S3 |
| 11) Evolutionary responses (LMMs) | MQ | Plant (nested in rep. pop.) | 2 N envs. × 2 host envs. | 14 rep. pops. × 1–4 plants | 47 | Fig. 3, SI Appendix Table S3 |
| 12) Retention of genetic variation (LMs) | P+ | Rep. pop. | 3 pop. types × 2 N envs. | 22 rep. pops. | 22 | Fig. 4, SI Appendix Table S4 |
| 13) Retention of genetic variation (LMs) | MQ | Rep. pop. | 2 N envs. × 2 host envs. | 14 rep. pops. | 14 | Fig. 4, SI Appendix Table S4 |
| 14) Plant growth over EE (LMMs) | All | Rep. pop. (repeated measures across GC) | 4 inoculation treatments × 2 N envs.× 4 GC | 8–20 rep. pops. per condition and GC | 392 | SI Appendix Table S5 |

pops. = population; rep. pop(s). = replicate population(s); envs. = environments; GC = growth cycles; dist. = distribution.

#### Supporting Figures

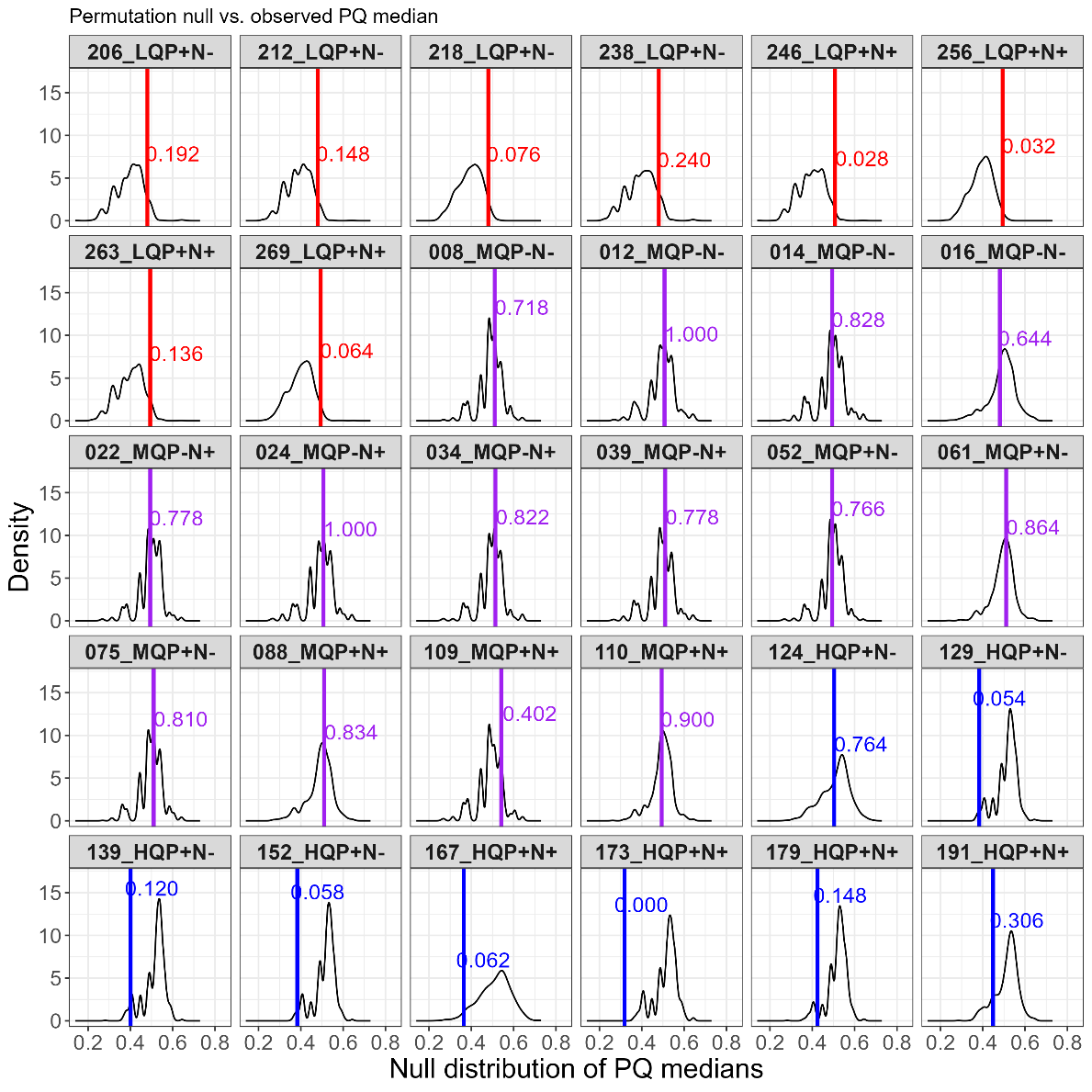

###### SI Appendix Figure S1: Permutation results assessing whether evolved populations differed from expectations under random sampling.

Permutation tests (1,000 iterations) accounting for replicate‑specific sampling effort are shown for each of the 30 independently evolved populations. For each replicate, the black distribution represents the null distribution of median partner quality (PQ) values generated by randomly drawing strains with replacement, with the number of draws matched to the number of nodules sampled for each replicate population. Coloured vertical lines indicate the observed median PQ of each evolved population, with blue, purple, and red corresponding to populations founded from the initially High‑, Medium‑, and Low‑Quality population types, respectively.

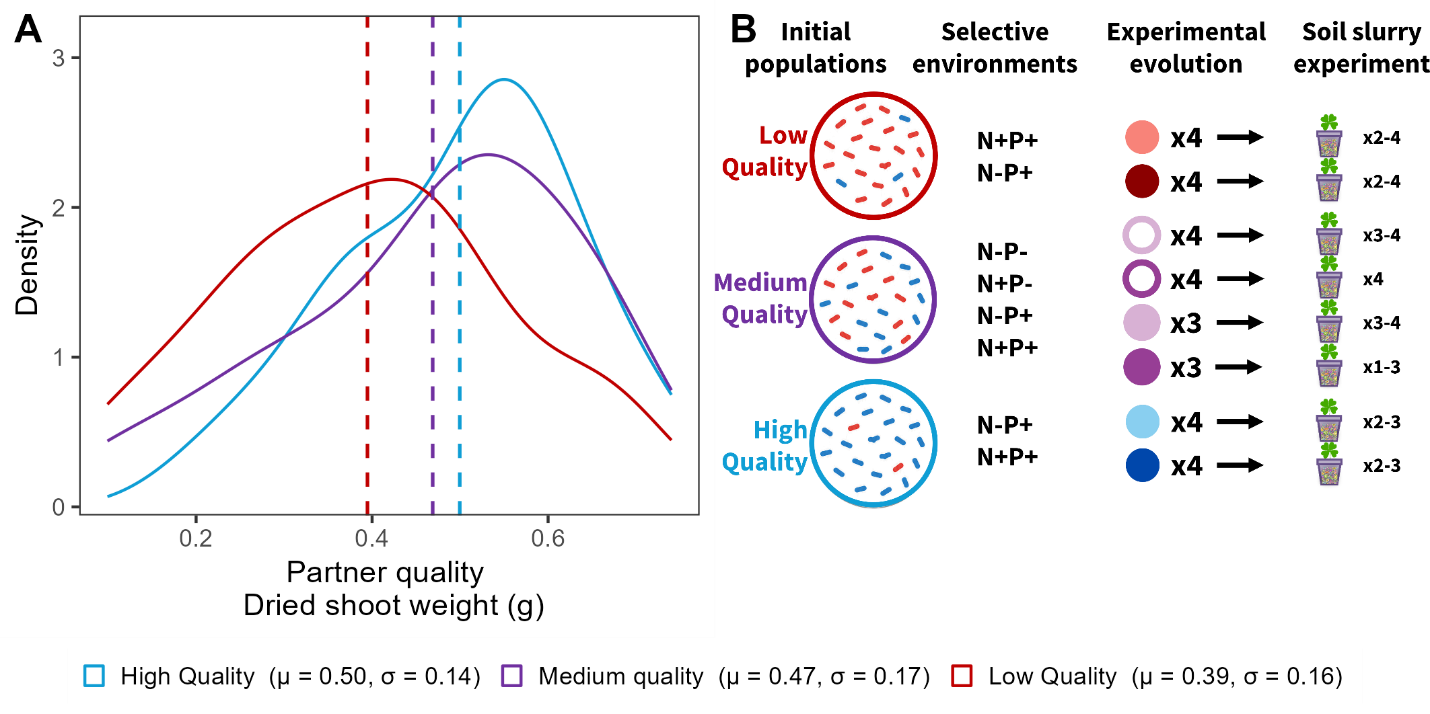

###### SI Appendix Figure S2: Initial partner quality distributions for each population type and experimental design overview.

**A** Kernel density plots of partner quality (dried shoot biomass) from the prior single‑strain assay (5). Coloured curves show the distribution for each population type; dashed lines mark mean values (*μ*). The legend reports each population’s mean (*μ*) and standard deviation (*σ*). **B** Overview of the experimental evolution design, including initial population types (N = 28 strains each), selective environments, number of replicate populations per population type and selective environment combinations, and number of plant replicates used in the soil‑slurry inoculation experiment.

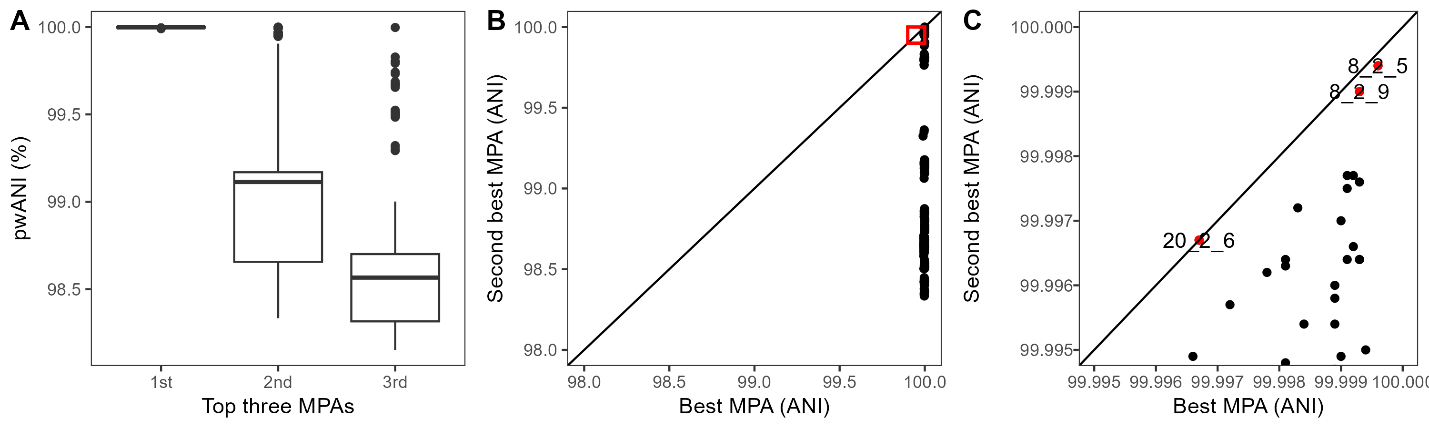

###### SI Appendix Figure S3: Performance of the ANI‑based match‑prediction approach to determine the most probable ancestor (MPA).

**A** Boxplots showing pairwise ANI values for the top, second‑best, and third‑best MPA hits. The sharp drop in ANI from the top hit to lower‑ranked hits indicates high specificity of the top match. **B** Pairwise ANI between each isolate and its top MPA hit (x‑axis) versus the second‑best hit (y‑axis). Most points fall below the 1:1 line, showing that second‑best hits are substantially less similar. **C** Zoomed‑in view of the ≥99.9% ANI region (red box in panel B). Three isolates approach the 1:1 line, indicating comparable similarity of their first and second hits. These isolates were resolved by assigning each to the MPA matching its original population type, based on their isolate IDs.

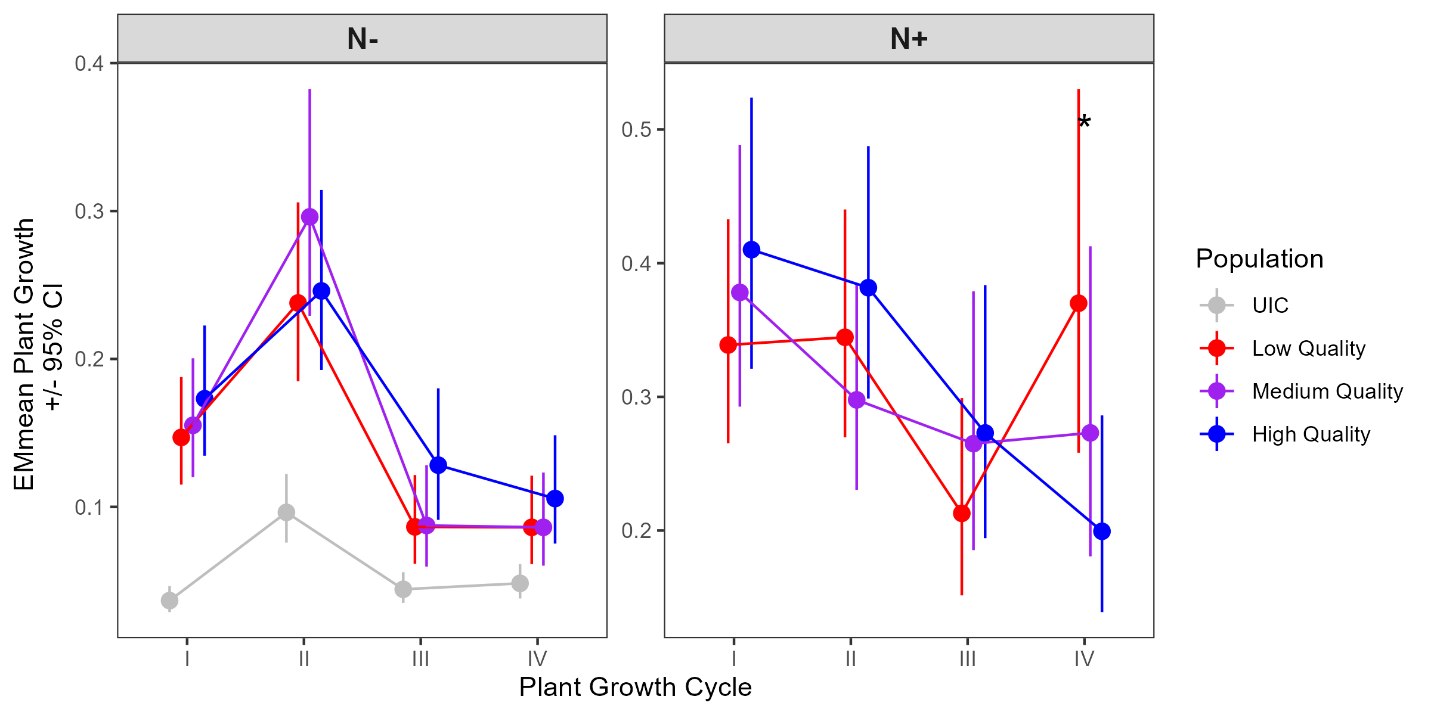

###### SI Appendix Figure S4: Time‑course of plant shoot biomass across four growth cycles in the experimental evolution study.

Points show estimated marginal means of dried shoot biomass from linear mixed-effects models (LMMs; see **SI Appendix Methods S3iv**), with 95% confidence limits. Analyses were based on repeated measurements of plants nested within independently evolved replicate populations across four sequential growth cycles. Panels compare nitrogen-limited (N−) and nitrogen-supplemented (N+) conditions. Colours indicate initial population type (High-Quality = blue, Medium-Quality = purple, Low-Quality = red), while gray lines/points represent uninoculated controls grown under N-free conditions. The asterisk in the N+ panel marks a significant difference between the Low- and High-Quality populations in the final growth cycle only.

#### Supporting Datasets

###### SI Appendix Dataset S1: Contrasts for linear mixed models assessing evolutionary responses to selective environments across different initial population types.

Statistical contrasts for shoot biomass, total nodule number, and per‑nodule weight from the soil slurry experiment. The dataset includes treatment contrasts comparing nitrogen environments (N-, N+) within each initial population type and host environment (P–, P+), as well as pairwise contrasts among population types within each nitrogen environment. Two model subsets are provided: Model I (P+ only) and Model II (Medium‑Quality population only). P‑values are adjusted using Tukey (pairwise) and Dunnett (treatment) corrections.

###### SI Appendix Dataset S2: Contrasts for linear models evaluating the effects of initial population type and selective environment on standing genetic variation.

The dataset includes pairwise contrasts among initial population types within selective environments, as well as treatment contrasts for nitrogen environments (within host environment) and host environments (within nitrogen environment). For each comparison, T statistics, P values, and effect sizes expressed as percent change with 95% confidence limits are reported.

###### SI Appendix Dataset S3: Metadata for the 363 isolates collected from nodules at the end of experimental evolution.

The dataset includes genome assembly statistics (completeness, contamination, total contig number, assembly length, GC content, and N50), taxonomic classifications generated using PGAP, the inferred most probable ancestor (MPA) for each isolate, and the experimental conditions under which each isolate was collected.

###### SI Appendix Dataset S4: Contrasts from linear mixed models evaluating the effects of initial population type, nitogen environment, and growth cycle on plant growth measured throughout experimental passaging.

The dataset includes pairwise contrasts among initial population types within both nitrogen treatments (N-, N+), as well as comparisons to uninoculated controls in the N– environment. For each contrast, model estimates include the test statistic, P value, and effect size expressed as percent change with 95% confidence limits.

#### SI References

1.	P. Somasegaran, *Handbook for rhizobia: methods in legume-Rhizobium technology* (Springer Science & Business Media).

13.	R Core Team, R: A Language and Environment for Statistical Computing. (2024). Deposited 2024.

14.	H. Wickham, RStudio, tidyverse: Easily Install and Load the “Tidyverse.” (2023). Deposited 22 February 2023.

15.	Y. Xie [aut, *et al.*, knitr: a general-purpose package for dynamic report generation in R. (2025). Deposited 20 December 2025.

16.	H. Zhu [aut, *et al.*, kableExtra: construct complex table with “kable” and pipe syntax. (2024). Deposited 24 January 2024.

17.	D. Bates, *et al.*, lme4: Linear Mixed-Effects Models using “Eigen” and S4. (2025). Deposited 2 December 2025.

18.	A. Kuznetsova, P. B. Brockhoff, R. H. B. Christensen, S. P. Jensen, lmerTest: tests in linear mixed effects models. (2026). Deposited 13 January 2026.
